# Altered chromatin occupancy of patient-associated H4 mutants misregulate neuronal differentiation

**DOI:** 10.1101/2023.09.29.560141

**Authors:** Lijuan Feng, Douglas Barrows, Liangwen Zhong, Kärt Mätlik, Elizabeth G. Porter, Annaelle M. Djomo, Iris Yau, Alexey A. Soshnev, Thomas S. Carroll, Duancheng Wen, Mary E. Hatten, Benjamin A. Garcia, C. David Allis

## Abstract

Chromatin is a crucial regulator of gene expression and tightly controls development across species. Mutations in only one copy of multiple histone genes were identified in children with developmental disorders characterized by microcephaly, but their mechanistic roles in development remain unclear. Here we focus on dominant mutations affecting histone H4 lysine 91. These H4K91 mutants form aberrant nuclear puncta at specific heterochromatin regions. Mechanistically, H4K91 mutants demonstrate enhanced binding to the histone variant H3.3, and ablation of H3.3 or the H3.3-specific chaperone DAXX diminishes the mutant localization to chromatin. Our functional studies demonstrate that H4K91 mutant expression increases chromatin accessibility, alters developmental gene expression through accelerating pro-neural differentiation, and causes reduced mouse brain size *in vivo*, reminiscent of the microcephaly phenotypes of patients. Together, our studies unveil a distinct molecular pathogenic mechanism from other known histone mutants, where H4K91 mutants misregulate cell fate during development through abnormal genomic localization.

## Introduction

Chromatin integrates intrinsic cellular and environmental cues to regulate diverse DNA-templated processes, including replication, transcription, and DNA damage repair. Mutations in chromatin regulators, including nucleosome remodelers, writers, readers, and erasers of histone and DNA modifications, have been emerging in cancer and developmental disorders^1,2^. Histones are the main proteins in chromatin, which together with DNA form nucleosomes and higher-ordered chromatin structures. Four core histone proteins, H2A, H2B, H3, and H4, are encoded by multiple paralogous genes each to meet the demand for histones arising during DNA replication in S-phase. Somatic missense mutations located in all four core histones were recently discovered in malignancies, which changed the chromatin signatures of cancer and highlighted the importance of histones in human disease^3–9^. Mechanistic studies of recurrent histone H3 mutations in pediatric cancers revealed that mutated histones can inhibit the enzymatic activity of the cognate histone methyltransferases, and lead to changes in the epigenetic landscape and gene expression^10–13^.

In addition to cancer-associated histone mutations, *de novo* germline mutations in genes encoding histone H4 and histone H3 variant H3.3 have recently been identified in 33 and 59 patients, respectively, with developmental disorders, suggesting histones are key regulators in diseases beyond cancer^14–20^. Patients bearing these *de novo* heterozygous mutations share a phenotype of developmental delay and intellectual disability. While several somatic histone mutations have been well studied in oncogenesis, the mechanistic and functional roles of these germline histone mutations in human development remain largely unknown. A recent study on germline mutations changing H3.3 glycine 34 to arginine (G34R) identified that H3.3 G34R decreases the methylation of the adjacent K36 residue and impairs recruitment of a DNA methyltransferase to alter the expression of immune and neuronal genes, which contribute to progressive microcephaly and neurodegeneration^20^. Almost all studies on disease-associated histone mutants to date have focused on histone H3.3, yet whether and how other germline histone mutations dysregulate gene expression and impact development is unclear.

Here our study focuses on mutations affecting lysine 91 of histone H4, which to date have been reported in seven patients presenting with microcephaly, intellectual disability, and dysmorphic facial features^14–16^. In all these cases, patients carry mutations in just one out of the fourteen H4 gene copies in the human genome, and these mutations change lysine 91 to either glutamine (Q), arginine (R), or glutamic acid (E) (**Figure S1A**). H4K91 is located in the globular domain of H4 and at the H2B-H4 dimer-tetramer interface of nucleosomes (**Figure S1B**). Consistent with this important structural localization within the nucleosome, mutated H4K91 destabilizes nucleosomes *in vitro*, as inferred by more active H2A/H2B dimer exchange^21^. Yet how mutations of the H4 histone fold impact development is not clear. In this study, we aim to characterize the binding properties of H4K91 mutants, their genomic localization, and their roles in regulating gene expression as well as brain development to reveal key underlying mechanisms of patient-associated H4 germline mutations. Our in-depth study on the dominant H4 mutants unveils that histone mutants act through abnormal genomic localization, which to our knowledge defines a novel molecular pathogenic mechanism compared to previous research that focused on H3.3 mutations affecting histone posttranslational modifications (PTMs). Furthermore, our study provides a compelling molecular connection between H4 mutants and the histone variant H3.3, contributing to the intricate regulatory landscape of the histone mutations in developmental disorders.

## Results

### H4K91 mutants display puncta formation and enhanced association with H3.3

The H4 protein is encoded by fourteen paralog genes, and heterozygous mutations of just one gene cause developmental defects in patients, suggesting that H4K91 mutants may have dominant effects on development. Therefore, we reasoned that expression of mutated H4 in the presence of endogenous wild-type H4 would recapitulate its functional effects, similar to effects observed with other histone mutations^10–13,22^. To characterize the function of mutated H4 during development, we first stably expressed HA-tagged wild-type H4 (H4WT) or H4K91 mutants in mESCs (**Figure S1C**). We focused subsequent studies on H4K91R and H4K91Q mutants as both are more prevalent in patients and more robustly accumulated in our model. Surprisingly, we found that mutated H4 appear as distinct nuclear puncta. In contrast, H4WT is homogeneously distributed within the nucleus (**Figure 1A**, **Figure S1D**). Quantification of the puncta intensity in cells with a wide range of expression levels confirmed that both H4 mutants, but not the wild-type H4, form distinct puncta (**Figure 1B**). Additionally, we observed the formation of discrete puncta in both live (**Video S1**) and fixed mESCs that transiently express GFP-tagged H4 mutants (**Figure S1E**), ruling out the possibility that the puncta identified by HA staining are caused by fixation and immunofluorescence staining^23^.

**Figure 1:**
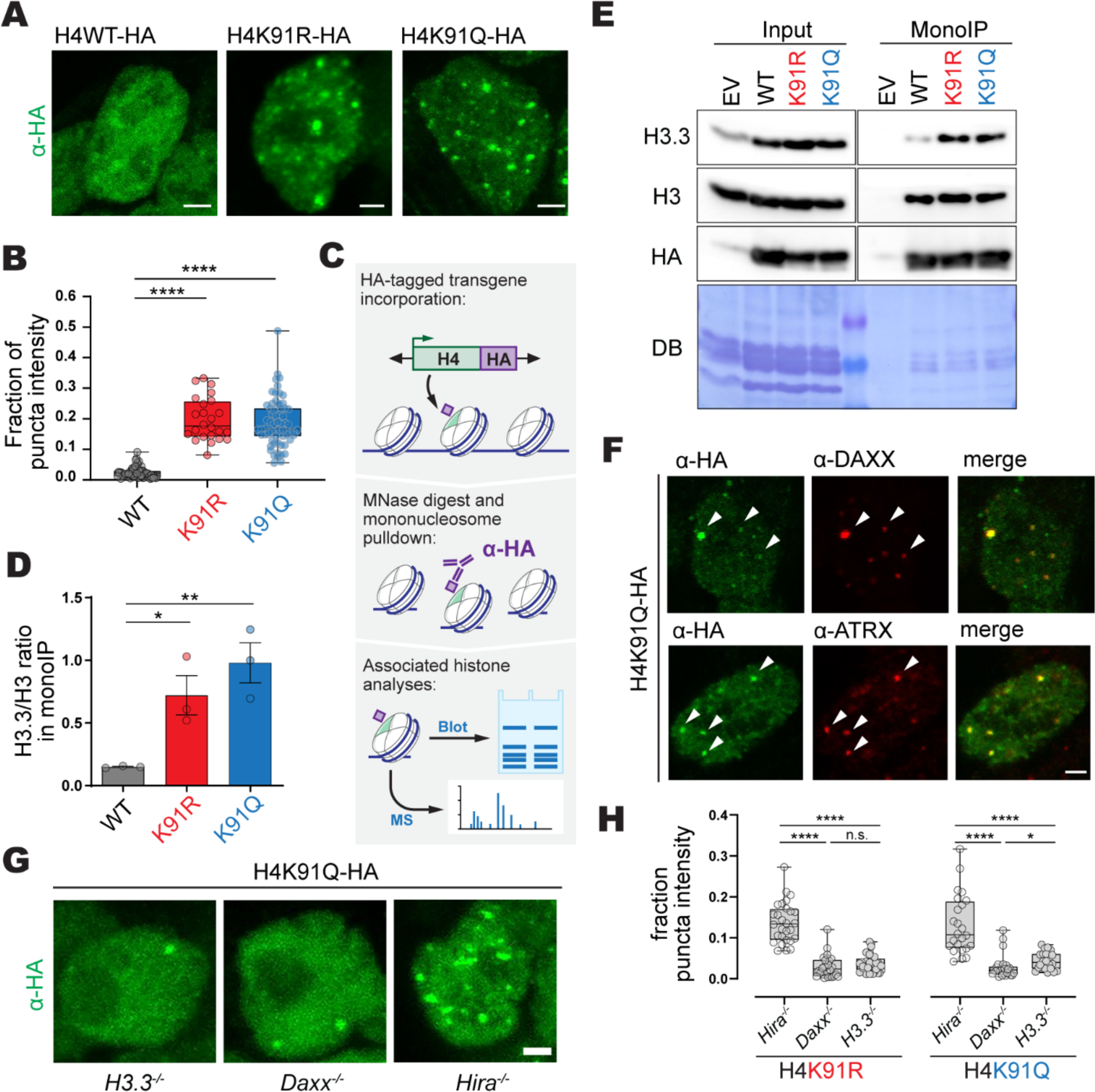
H4K91 mutants enhance association with the histone variant H3.3 and form distinct nuclear puncta. (**A**) Both H4K91 mutants display a puncta pattern. Immunofluorescence images of mESCs that express HA-tagged H4WT or H4K91 mutants (K91R or K91Q) with HA antibody. Images are Z-stack projections. Scale bar: 2μm. (**B**) Quantification of in-puncta fluorescence intensity in the nucleus. Fluorescence intensity within the puncta is divided by the whole nucleus intensity and displayed as a box plot. The box plot shows all data points with the median, and the boundaries indicate the 25^th^ and the 75^th^ percentiles. Each dot represents one cell. Statistical significance was calculated using an unpaired t-test. **** *p*<0.0001. (**C**) Schematic of mononucleosome IP method. Chromatin from 293T cells expressing WT or H4K91 mutants are first subject to MNase digestion. Then, mononucleosomes containing HA-tagged H4WT or H4K91 mutants are pulled down with HA antibody and analyzed by mass spectrometry (MS) analysis or western blot (WB). (**D**) H4K91 mutants display increased binding with the histone variant H3.3. A plot of the H3.3/H3 ratio based on MS analysis of the pulled-down nucleosomes. Three independent replicates were done. Error bars represent mean ± s.e.m, and statistical significance was calculated using an unpaired t-test. * p<0.05, ** p<0.005. (**E**) WB of the pulled-down mononucleosomes with the following antibodies: HA, H3.3 specific antibody, and H3 (which recognizes both canonical histone H3 and variant H3.3). (**F**) H4K91 mutant cells staining with the H3.3-specific chaperones DAXX and ATRX. Immunofluorescence images of mESCs expressing a HA-tagged H4K91 mutant. The following antibodies were used: HA, DAXX, and ATRX. Arrowheads indicate colocalization. A single Z stack image is shown. Scale bar: 2μm. (**G**) Immunofluorescence images of H3.3−/−, DAXX −/− and Hira−/− mESCs that express HA-tagged H4K91Q. Scale bar: 2μm. (**H**) Quantification of in-puncta HA fluorescence intensity in different mESCs. Each dot represents one cell. Statistical significance was calculated using an unpaired t-test. * *p*<0.05. **** *p*<0.0001. See also Figure S1.

To understand the puncta formation of mutated H4, we sought to characterize its chromatin deposition pathway. During nucleosome assembly, H4 is deposited onto DNA within the H3/H4 tetramer, followed by the association of two H2A/H2B dimers. H4 is the only invariant core histone and thus binds to both canonical H3, deposited by CAF1, and the specialized histone variant H3.3, deposited in euchromatin regions by Hira, and in repetitive heterochromatic regions by ATRX and DAXX chaperones^24–28^. Because of the fundamental importance of the H3/H4 interaction in forming the nucleosome, we first asked whether H4K91 mutants can bind to both H3 and H3.3. We tested this by mononucleosome immunoprecipitation using HEK293T cells expressing HA-tagged H4WT or H4K91 mutants, followed by mass spectrometry. We found that H4K91 mutants preferentially associate with H3.3 over canonical H3, displaying a robust five-fold increase compared to WT H4 (**Figure 1C-D**). These results were further validated by western blotting with an H3.3-specific antibody (**Figure 1E**). Previous studies characterized H3.3 puncta in mESCs arising from deposition into heterochromatic regions, including telomeres ^29–31^. Consistent with increased binding to H3.3, H4K91 mutant puncta colocalized with the telomere probe and mutated H4-containing nucleosomes carried higher K9-trimethylated histone H3 (H3K9me3), a canonical heterochromatin mark (**Figure S1E, F**).

As the heterochromatin deposition of H3.3 is facilitated by the chaperone protein ATRX and DAXX, we investigated the localization of ATRX and DAXX in mESCs expressing mutated H4^25,31^. Notably, these H4K91 mutant nuclear puncta co-localize with DAXX and ATRX (**Figure 1F**). To probe whether histone variant H3.3 and these chaperones are essential for H4 mutant puncta formation, we, expressed mutated H4 in mESC lines lacking H3.3, DAXX, or Hira, and analyzed H4 mutant localization. Staining and quantification of the puncta revealed that depletion of H3.3 or DAXX, but not the euchromatin-specific H3.3 chaperone HIRA, abolished the H4K91 mutant puncta formation in mESCs (**Figure 1G, H**). Altogether, these data show that H4K91 mutants are preferentially associated with H3.3 and deposited into heterochromatin by a DAXX/ATRX-dependent pathway.

### H4K91 mutants are deposited into H3.3 and H3K9me3-enriched heterochromatin

DAXX and ATRX cooperate to deposit the H3.3/H4 complex into several distinct heterochromatic regions, including telomeres, pericentromeric regions, and interspersed repeats^31–34^. To define the incorporation sites of mutated H4 in mESCs, we profiled the genome-wide distribution of mutated H4K91 using chromatin immunoprecipitation followed by high-throughput DNA sequencing (ChIP-seq). After clustering H4K91 peaks based on the H4 ChIP signal, we found that both H4K91 mutants occupied a shared subset of new genomic loci (referred to as “K91 mutant-enriched peaks”), compared to H4WT. Interestingly, while H4K91Q localization was primarily confined to these shared mutant-enriched peaks, H4K91R was also localized to additional areas that also contain H4WT (referred to as “other peaks”), suggesting a wider deposition of the arginine mutant (**Figure S2A**). Further annotation of the shared K91 mutant-enriched peaks revealed that the majority (55%) of these peaks localized at distal intergenic regions, whereas most (52%) other peaks are at promoter regions (**Figure S2B**). We reasoned that the disease-causing function of H4 mutants is mediated by ectopic localization to *de novo* occupied regions shared across all mutants, represented by K91 mutant-enriched peaks. A closer examination of these peaks found that they have abundant H3.3 signal as well as H3K9me3, a marker of heterochromatin (**Figure 2A**, **Table S1**). Using the regions under both H4WT and H4K91 mutant peaks, we found that H4K91 mutant ChIP signal had a strikingly higher correlation with H3.3 and H3K9me3 ChIP signals than H4WT (**Figure 2B**), further corroborating the enhanced binding of mutant H4 to H3.3 documented in the pulldown assay (**Figure 1D, E**). Notably, K91 mutant-enriched peaks showed enrichment at several selected targets of H3.3 and H3K9me3, which were identified previously in mESCs (**Figure 2C**)^32,33^.

**Figure 2:**
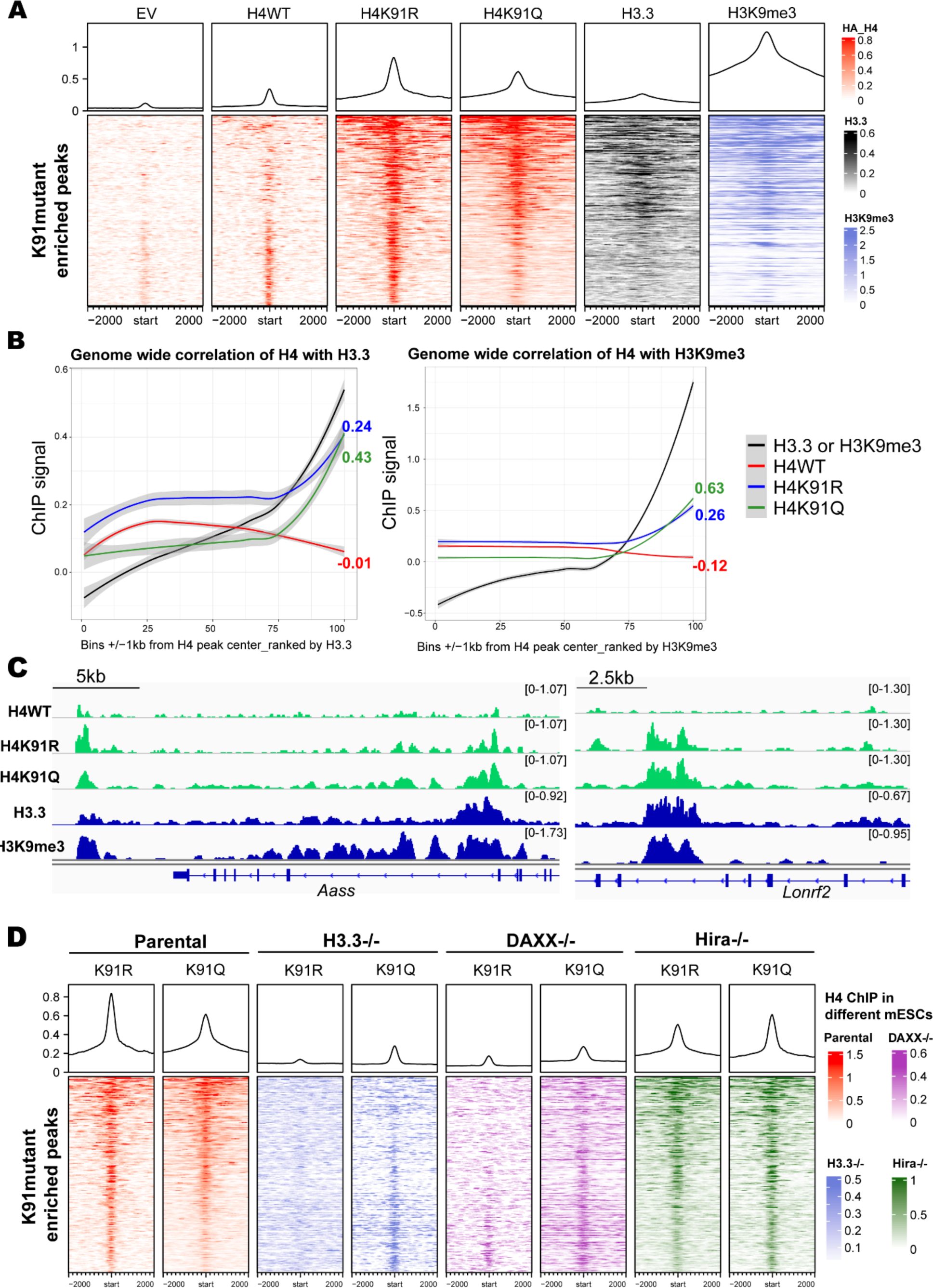
H4K91 mutants are recruited to H3.3 and H3K9me3 enriched heterochromatin regions. (**A**) Heatmap representation of histone H4 bound chromatin peaks that have increased occupancy by H4K91 mutants (H4K91R and H4K91Q) compared to H4WT in mESCs (Table S1). After removing peaks in cells expressing empty vector (EV), peaks from H4WT, H4K91R, and H4K91Q expressing cells were clustered based on the HA signal. The cluster containing H4K91 mutant enriched peaks is shown, together with the ChIP-seq signal of histone H3 variant H3.3 and H3K9me3. These regions represent ± 2kb from the center of the H4 peaks. The color scheme represents the ChIP-seq signal density with darker indicating increased signals. (**B**) Correlation of H4WT or H4K91 mutant ChIP-seq signal with H3.3 (left) and H3K9me3 (right) ChIP-seq signal within H4 peak regions in mESCs. Input normalized ChIP-seq signal within bins that comprise regions ±-1kb around the center of H4 peaks (union of H4WT, H4K91R, and H4K91Q) was used for this Pearson’s correlation coefficient (PCC) analysis, and PCC values are shown to the right of each figure. (**C**) Genome browser representative tracks of HA, H3.3, and H3K9me3 ChIP–seq signals at selected H4 mutant target genes in mESCs. (**D**) Heatmap of H4K91 mutant enriched peaks in Parental, H3.3−/−, DAXX−/−, and Hira −/− mESCs expressing H4K91R-HA or H4K91Q-HA. See also Figure S2, S3 and Table S1.

Next, we asked if the loss of H3.3 or its specific chaperone DAXX would blunt ectopic H4 enrichment, similar to the rescue of puncta in the knockout (KO) mESCs (**Figure 1G, H**). We found that K91 mutant-enriched peaks were diminished in H3.3 or DAXX KO mESCs but are maintained in Hira KO mESCs (**Figure 2D**). This is confirmed by the loss of genome-wide correlation of H4K91R or H4K91Q occupancy with H3.3-enriched sites (**Figure S2C**). Collectively, these data suggest that H4K91 mutants are incorporated into distinct genomic loci enriched in H3.3 and H3K9me3, and this ectopic recruitment relies on the histone variant H3.3 and its chaperone DAXX.

The observation of distinct puncta in the nucleus and distinct H3.3-dependent genomic localization of mutated H4 prompted us to study the potential effects of mutant expression on chromatin structure. Lysine 91 of H4 localizes at the H2B/H4 dimer tetramer interface, and this residue forms a salt bridge with an E76 residue in histone H2B (**Figure S1B)**. H4K91 mutants are known to destabilize the nucleosome *in vitro* and in yeast^21,35^. Moreover, histone mutants affecting another H2B/H4 dimer-tetramer interface residue, H2B E76K, have been reported to destabilize nucleosomes *in vitro* and significantly promote chromatin accessibility^8,9,21^. To test whether disease-associated H4K91R/Q mutants alter chromatin accessibility in mammalian cells, we visualized accessible DNA using a previously reported assay of transposase-accessible chromatin with visualization (ATAC-see)^36^. Using the HDAC inhibitor Panobinostat as a positive control, we observed foci of increased accessibility in HEK293T cells that express K91 mutants, but not wild type control (**Figure S3A-C**). Altogether, we found that expression of patient-associated H4K91 mutants increases chromatin accessibility in mammalian cells.

### H4K91 mutants alter mammalian brain development

We next investigated how the incorporation of H4K91 mutants could misregulate development. Patients carrying H4K91 mutations present with microcephaly, likewise an ectopic expression of H4K91 mutants (K91R/Q) in zebrafish led to underdeveloped brain^15^, together suggesting that a fraction of mutated H4 is sufficient to affect brain development *in vivo*. We sought to investigate the effect of H4K91 mutants on brain development in a mammalian system while minimizing the effects of genetic background, age, and sex. To achieve this, we generated H4WT and H4K91 mutant isogenic all-ESC mice using a tetraploid complementation approach. Specifically, we injected mESCs expressing HA-tagged H4 into tetraploid blastocysts, where the 4n cells of the host embryo contribute solely to the placenta, while the injected ESCs form the embryo proper^37^. H4K91 mutant pups at postnatal day 0 (P0) exhibited a reduction in brain size compared to H4WT mice, with no significant changes in body weight (**Figure 3A**). This phenotype mimics the microcephaly phenotypes observed in patients, suggesting that mice are a suitable model to study the underlying molecular mechanisms of H4K91 mutations. To visualize the localization pattern of WT and mutated H4 *in vivo*, we stained cerebral cortex sections with anti-HA antibody. Remarkably, H4 mutants show a similar pattern of nuclear puncta, and the puncta colocalize with DAPI-bright heterochromatin regions (**Figure 3B**). These findings recapitulate the distinct puncta detected in mESCs (**Figure 1A, F**, **Figure S1D-E**), suggesting that H4K91 mutants are consistently incorporated into heterochromatin in differentiated neurons *in vivo*.

**Figure 3:**
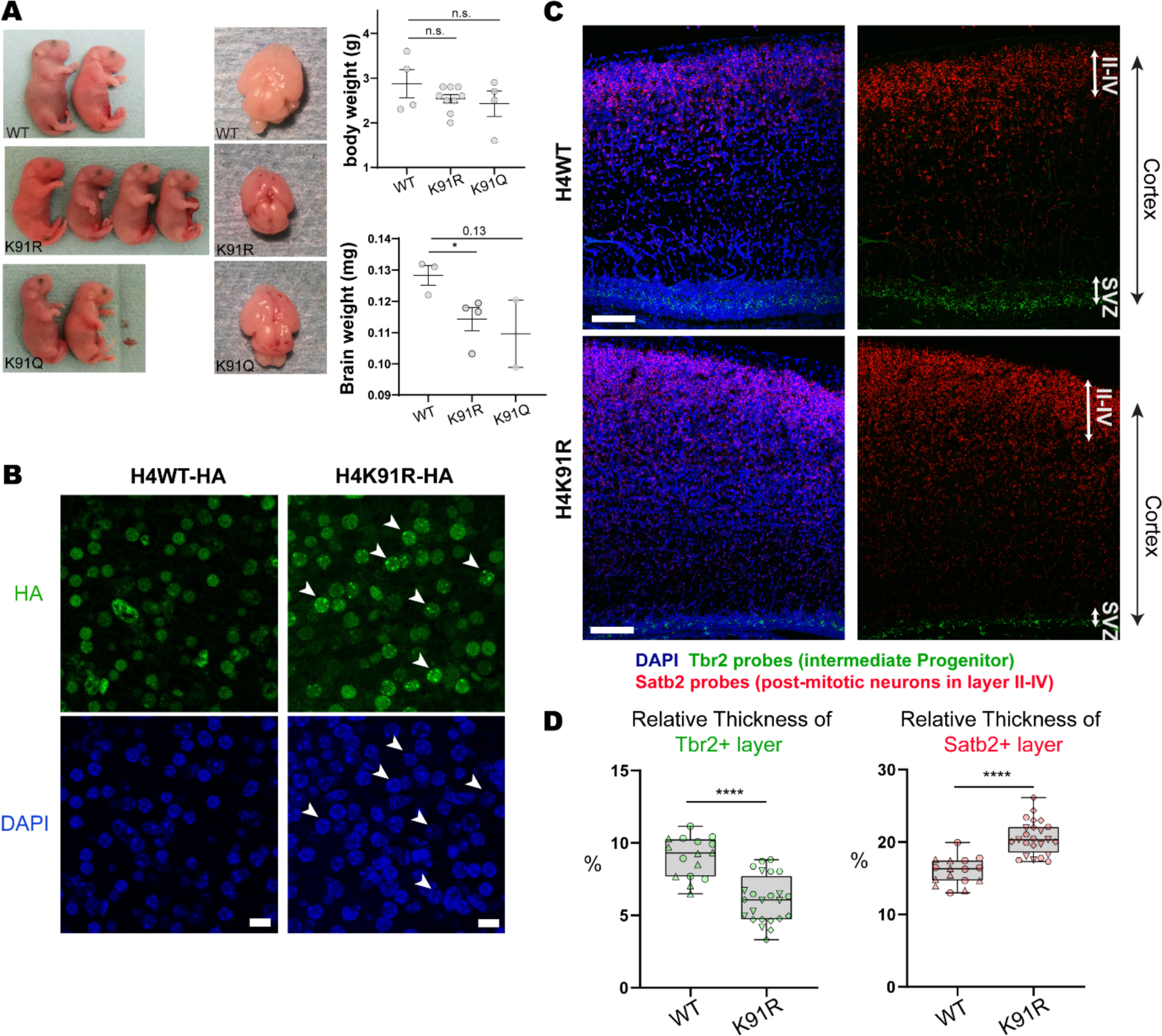
H4K91 mutants affect mouse brain development. (**A**) Left: Representative images of pups and brains of H4WT and H4K91 mutant isogenic mice at P0, which were generated through tetraploid complementation. Right: Quantification of body and brain weight. Box plots show mean ± s.e.m., and p-values are calculated by unpaired t-test. * *p*<0.05 (**B**) Representative immunofluorescence staining of mouse brain cerebral cortex with HA and DAPI. H4K91R shows puncta (arrowheads) pattern, which is also in DAPI bright regions. Scale bar: 10μm. (**C**) *In situ* hybridization of mice brain cerebral cortex with probes targeting transcripts of intermediate progenitor marker gene *Tbr2* (green) and post-mitotic neuron marker *Satb2* (red). Tbr2-positive intermediate progenitor cells localize in the subvertical zone (SVZ) of the cortex, while Satb2-positive post-mitotic neurons are in the upper layer of the cortex (II-IV). The full cortex is indicated by black arrows. Scale bar: 100μm. (**D**) Quantification of the relative thickness of Tbr2 and Satb2 positive layers (thickness of Tbr2/Satb2 layer divided by the thickness of the whole cortex). 8 different section regions are quantified for each mouse (n=2 H4WT-HA and n=3 H4K91R-HA). The box plot shows all data points with the median, and the boundaries indicate the 25^th^ percentile and 75^th^ percentiles. **** *p*<0.0001. p-values are calculated by unpaired t-test.

In the cerebral cortex, neuronal progenitor cells localize to the ventricular and subventricular zone (SVZ), and undergo proliferation, differentiation, and migration to give rise to neurons and glial cells, providing a system to study lineage specification and cell differentiation^38^. To investigate the effect of H4K91 mutant expression in neurogenesis, we performed *in situ* hybridization with two probes targeting intermediate progenitor cells (*Tbr2*) and post-mitotic neurons (*Satb2*). In both wild type control and mutant cerebral cortex, we observed *Tbr2* labeled progenitor cells localized at SVZ, while *Satb2* labels cells in the upper cortical layers (**Figure 3C**). We quantified the relative thickness of *Tbr2* or *Satb2* positive cells normalized to full cortex thickness in different section areas from independent mice. These analyses revealed that the H4 mutant brain has a thinner *Tbr2+* layer and expanded *Satb2+* layer compared to H4WT (**Figure 3D**). These changes indicate that mutated H4 misregulates neurogenesis *in vivo*, resulting in a reduced progenitor population.

### Accelerated neural differentiation occurs upon expression of H4K91 mutants

The H4K91 mutant-dependent phenotypes observed in mice mirror those of H3.3 loss, which promotes premature neural differentiation and altered gene expression in postmitotic neurons^39,40^. Since H4K91 mutants are recruited to H3.3-enriched genomic regions and are preferentially associated with H3.3 (**Figure 1D, E**, **Figure 2A-B**), we reasoned that H4 mutants, similar to the loss of H3.3, might misregulate neuronal differentiation and alter gene expression in differentiated neural cells. Therefore, we next investigated the effects of H4 mutants on neural differentiation. We differentiated mESCs to neural cells and detected Sox2+ NPCs and Map2+ neuron cells by immunostaining, indicating successful differentiation into the neuronal lineage (**Figure S4A-B**). We next asked whether H4 mutant expression alters gene expression by comparing transcriptional programs of four-day differentiated neural cells that express H4WT or mutant H4. We found overlapping transcriptional changes caused by H4K91R and H4K91Q mutants, further demonstrating that H4K91 mutants share the same mechanism (**Figure 4A**). Corroborating our findings from the developmental model, expression of H4K91R or H4K91Q caused upregulation of genes enriched in mature neuronal pathways, including synapse organization, neuron projection guidance and neuron migration (**Figure 4B**, **Table S2**), including known neuronal markers *Map2*, *Tubb3*, *Rbfox3*, and *Dcx*. (**Figure 4C**). Collectively, we demonstrate that H4K91 mutants alter gene expression and impact neural differentiation.

**Figure 4:**
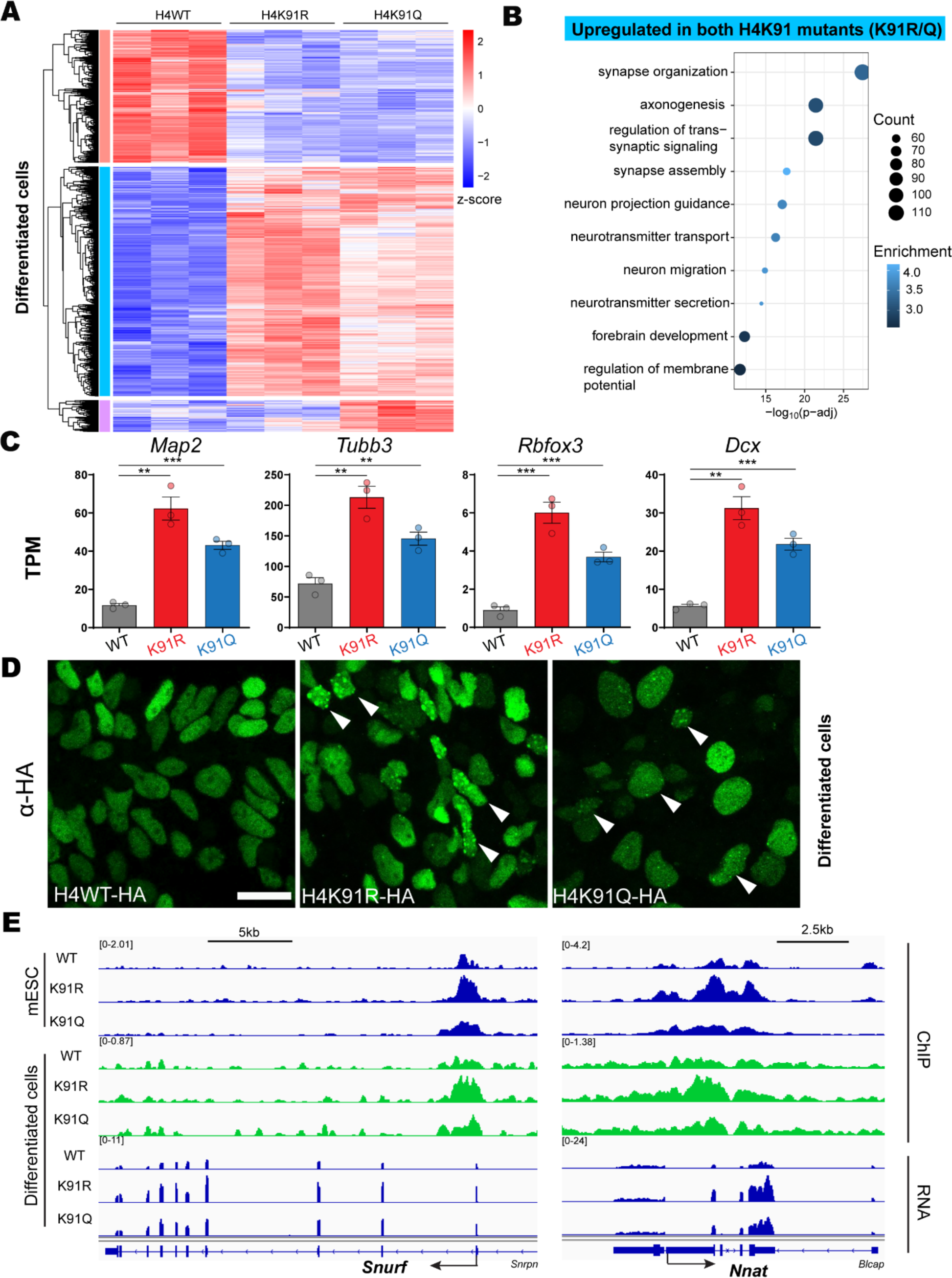
H4K91 mutants accelerate neural differentiation and alter gene expression. (**A**) Clustered heatmap representation of genes that are differentially expressed (with a fold change of 2 or more, and *p*<0.05) in neural cells after four days of differentiation that express WT or mutant H4. The color indicates the z-score of expression values for each gene. (**B**) Gene Ontology (GO) analysis of the upregulated cluster of genes (from Figure 4A) that are common to K91R and K91Q in differentiated neural cells (n=1240 genes; Table S2). (**C**) Visualization of mRNA expression of several mature neuron genes at four-day neural differentiation. The data represents mean ± s.e.m from three independent replicates. Statistical significance was calculated using an unpaired t-test. ** *p*<0.01. *** *p*<0.001. (**D**) Representative immunofluorescence images of four-day differentiated neural cells stained with HA antibody. Arrowhead labels point to cells with puncta patterns of H4K91 mutants. Scale bar: 20μm. (**E**) Genome browser representation of HA ChIP–seq signals for H4 in mESCs and differentiated cells, as well as the representative RNA-seq tracks at selected imprinted genes in differentiated cells. See also Figure S4 and Table S2.

We next investigated the nuclear localization and the chromatin occupancy of H4K91 mutants in differentiated neural cells. Similar to mESCs (**Figure 1A, F**, **Figure S1D-E**) and neural cells in the brain (**Figure 3B**), staining with anti-HA antibody identified the distinct puncta staining pattern in differentiated neural cells that express H4 mutants, but not WT H4 (**Figure 4D**). This implies that H4K91 mutants share the ectopic recruitment pattern in distinct cell lineages. We further performed anti-HA ChIP-seq to characterize the genome-wide incorporation of H4 mutants. Previous studies have shown that most co-enrichment of H3.3 and H3K9me3 in mESCs is lost upon differentiation to NPCs, except at telomeres^32,33^. Similarly, we saw enhanced chromatin occupancy of H4K91 mutants at telomeric regions in NPCs or differentiated neural cells (**Figure S4C**). Additionally, H3.3 and H3K9me3 have previously been shown to localize at imprinted genes, which are specialized genes that are expressed in a monoallelic fashion^34^. We focused on 20 known imprinted loci that were shown to carry H3K9me3^41^. We found the enrichment of H4K91 mutants at these imprinted loci in both mESCs and differentiated cells (**Figure S4D**). Interestingly, 12 out of 26 genes within these imprinted loci with enhanced occupancy of H4 mutants were significantly upregulated in mutant expressing neural cells (*p*<0.05, **Figure S4D)**, as exemplified by select gene loci (**Figure 4E**). Taken together, the distinct puncta staining, combined with the increased enrichment of H4K91 mutants at telomeres and imprinted genes, suggest that the same co-localization with H3.3 and H3K9me3 identified in mESCs is also occurring in differentiated neural cells.

## Discussion

Our study details how dominant histone H4 mutants at lysine 91 act through abnormal genomic localization, which to our knowledge demonstrates a new molecular pathogenic mechanism compared to previous research that focused on histone PTMs. We show that histone H4K91 mutants found in developmental disorders are incorporated into chromatin and preferentially associate with the histone variant H3.3, a histone variant that also has mutations in developmental disorders. Mediated by the H3.3-specific chaperone DAXX, the H3.3/H4K91 mutant complex is deposited into H3K9me3-enriched heterochromatin and forms distinct puncta in the nucleus of several distinct cell lineages (**Figure S5**). Expression of H4K91 mutant drives increased chromatin accessibility, consistent with our previous finding that H4 lysine 91 is critical for nucleosome stability *in vitro*^21^. Moreover, the accumulation of H4K91 mutants alters gene expression and promotes precocious neural differentiation, consistent with the reduced brain size and altered cortical layers observed in our mouse model (**Figure S5**). These findings recapitulate the microcephaly phenotype of patients carrying these germline mutations and are in line with the previously reported underdeveloped brain phenotype in H4K91 mutant zebrafish^15^. Our model will be instrumental in future efforts to elucidate the brain defects at distinct developmental stages, the cell-type specific genomic incorporation of H4K91 mutants into chromatin, as well as changes in chromatin accessibility and gene expression in post-mitotic neurons *in vivo* at distinct development times. These studies will clarify how H4K91 mutants lead to brain defects during development.

Like many histone lysine residues, H4K91 is subject to several post-translational modifications, including acetylation, mono-ubiquitination, and glutarylation^35,42,43^. A primary future direction will be to characterize the genome-wide distribution of H4K91 modifications in mutant expressing cells. So far, three different missense mutations have been identified in patients: H4K91E, H4K91Q, and H4K91R, which mimic glutarylated, acetylated, and unmodified lysine, respectively. All patients share similar clinical features, such as microcephaly and delayed development, which suggests that three mutated residues (E/R/Q) of H4K91 share a common mechanism of action in driving developmental phenotype. We propose that despite mimicking different post-translational modifications, these three documented H4K91 mutants converge to affect chromatin assembly and structure through nucleosome destabilization, altering basic cellular functions, such as DNA damage response and gene transcription, during development.

Besides histone H4K91, several *de novo* missense germline mutations affecting multiple residues of histone variant H3.3 have been reported in patients with progressive neurologic dysfunction and congenital anomalies^17–20^. A recent study of H3.3G34R substitution in mice found that the G34R mutant led to progressive microcephaly and neurodegeneration through impairing recruitment of the DNA methyltransferase DNMT3A and its redistribution on chromatin^20^. This study provides insights into the impact of H3.3 mutants on the chromatin landscape and highlights the roles of H3.3WT in regulating brain function. Furthermore, our study demonstrates that there is an enhanced association of H4K91 mutants with H3.3 and genome-wide colocalization of H4K91 mutants with H3.3. This study provides an intriguing molecular link between H4K91 mutants and the histone variant H3.3. We propose that H4K91 mutants and some H3.3 mutations may be incorporated into similar genomic regions, disrupting the local chromatin structure and altering gene expression. Further studies using H4K91 mutant mice and H3.3 conditional KO mice may help clarify the underlying regulatory mechanisms in the physiologic context of development. In light of the increasing number of histone germline mutations and the lack of understanding of their impact, our study here provides interesting biochemical, cellular, and molecular evidence for how mutations of H4 impact its binding properties with histone H3, altering H4 genomic incorporation, as well as derailing gene expression and cell fate during neural differentiation.

In addition to newly identified histone mutants, disturbances of imprinted gene expression due to genetic and epigenetic defects account for imprinting disorders, a group of congenital diseases affecting growth and development^44^. The enrichment of H4K91 mutants at imprinted loci and upregulation of several imprinted genes with H4K91 mutant expression suggests that mutated H4 might cause developmental defects through the aberrant expression of imprinted genes, which is one focus of our future studies. Together, our findings shed light on the effects of development by histone mutants beyond their well-known oncogenic roles.

## Acknowledgments

We thank past and present members of the Allis laboratory, especially Leah Gates, Agata Lemiesz Patriotis, and Marylene Leboeuf. We acknowledge The Rockefeller University for financial support and the community, especially Shixin Liu, Viviana Risca, Hironori Funabiki, and Michael W. Young for helpful discussions and support. We thank Shasha Chong for sharing the ImageJ algorithm used to quantify the puncta intensity. We thank Francisca Vitorino from Garcia lab for uploading the Mass Spectrometry data to the MassIVE database. Finally, we acknowledge the help from the following resource centers at The Rockefeller University: Bio-Imaging, Bioinformatics, and Genomics. This work was supported by the NIH grant P01CA196539 to C.D.A. L.F. was supported by the C. H. Li Memorial Scholar Fund at The Rockefeller University and an NIH Career Award (K99HD107908). E.G.P. and B.A.G. were supported by NIH grants R01NS111997 and R01HD106051. L.Z. and D.W. were supported by the NIH grant (R01GM129380 to D.W.). A.A.S. is supported by UTSA and NCI grant R01CA234561. This article is dedicated to the memory of C. David Allis, who passed away on January 8, 2023.

## Author contributions

L.F. and C.D.A. conceptualized and initiated this project. L.F. performed most experiments with the help of L.Z. and K.M. to generate the mice and collect the brain samples. D.B. conducted all bioinformatic analyses. E.G.P. performed mass spectrometry and data analysis. A.D. and I.Y. contributed to the cell culture assays. A.A.S. provided conceptual advice and designed the cartoon figure. T.S.C, D.W, M.E.H, B.A.G., and C.D.A. participated in study supervision. L.F. wrote the manuscript with the input from all authors.

## Data availability

The ChIP-seq and RNA-seq data have been deposited to the Gene Expression Omnibus (GEO) database under accession number GSE231567. The mass spectrometry data has been deposited to the MassIVE database under dataset number MSV000091837. Any additional information or materials are available upon request.

---

**Figure S1:**
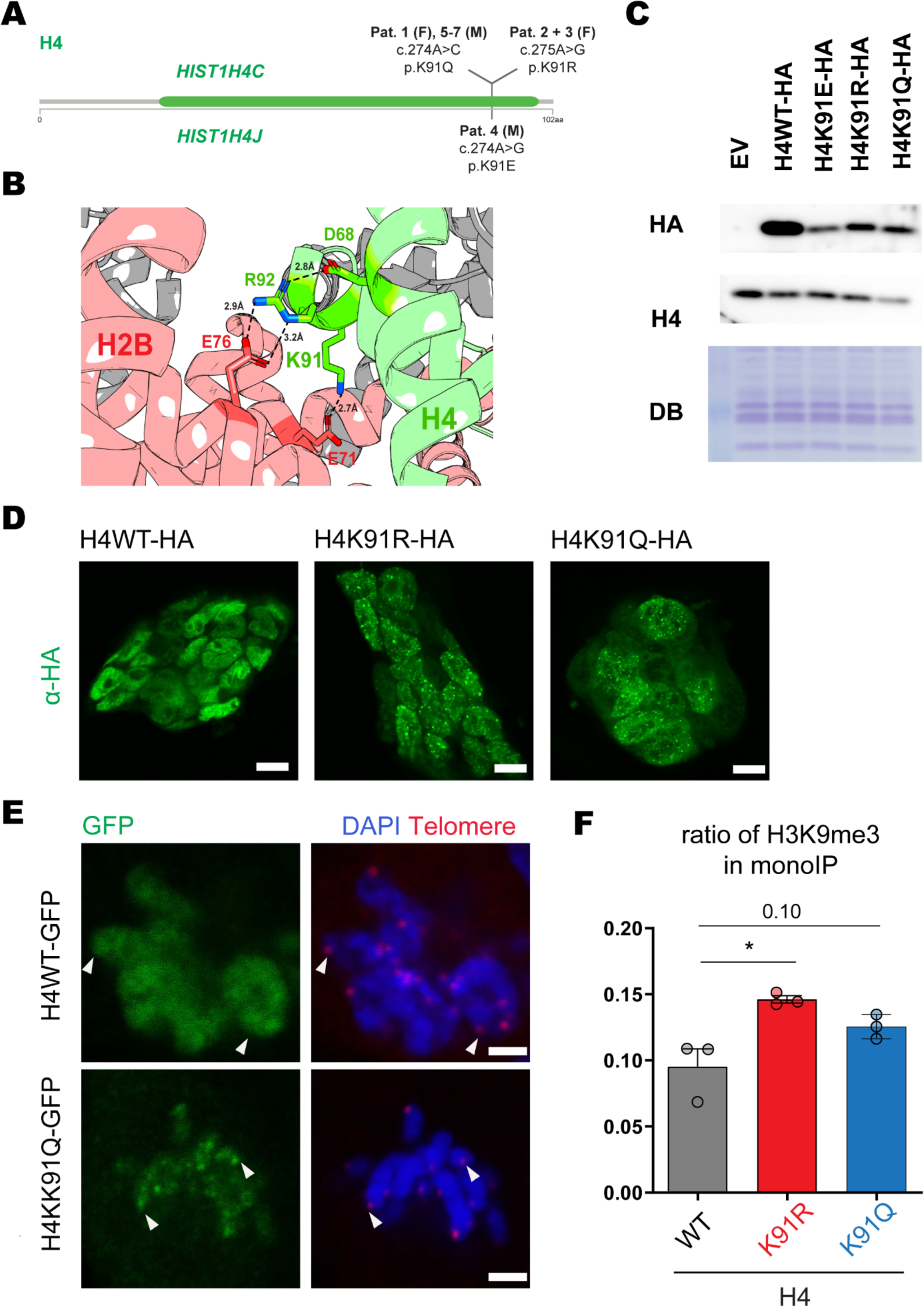
H4K91 mutants associated with developmental disorders form distinct puncta and colocalize with telomeres in mESCs. Related to Figure 1. (**A**) Summary of seven patients carrying germline mutations affecting lysine 91 of histone H4. The globular domain of H4 is indicated by the green bar. Two genes encoding the H4 protein are affected: HIST1H4C and HIST1H4J. The HGVS nomenclature for each substitution is labeled. F: female, and M: male. (**B**) H4K91 is localized at the H2B–H4 dimer-tetramer interface. H4K91 can form a salt bridge with H2BE71, while H4D68 and H4R92 form hydrogen bonding with H2BE76. (**C**) Western blot analysis of mESCs that express HA-tagged H4WT and H4K91 mutants (K91E, K91R, or K91Q). HA staining detects the expression level of the transgene. The expression level is less than the endogenous level and is not detected by anti-H4 staining. (**D**) Immunofluorescence images of mESCs colonies that express HA-tagged H4WT or H4K91 mutants. Scale bar: 10μm. (**E**) In situ hybridization of mESCs that transiently express GFP tagged WT or H4K91Q with telomere probe (TelGCy3). Arrowheads represent the telomeric regions. Single Z stack images of mitotic cells are shown. Scale bar: 2μm. (**F**) A plot of the H3K9me3 ratio based on MS analysis of the pulled-down mononucleosome in Figure 1C. The data represents mean ± s.e.m from three independent replicates. Statistical significance was calculated using an unpaired t-test. * *p*<0.05.

**Figure S2:**
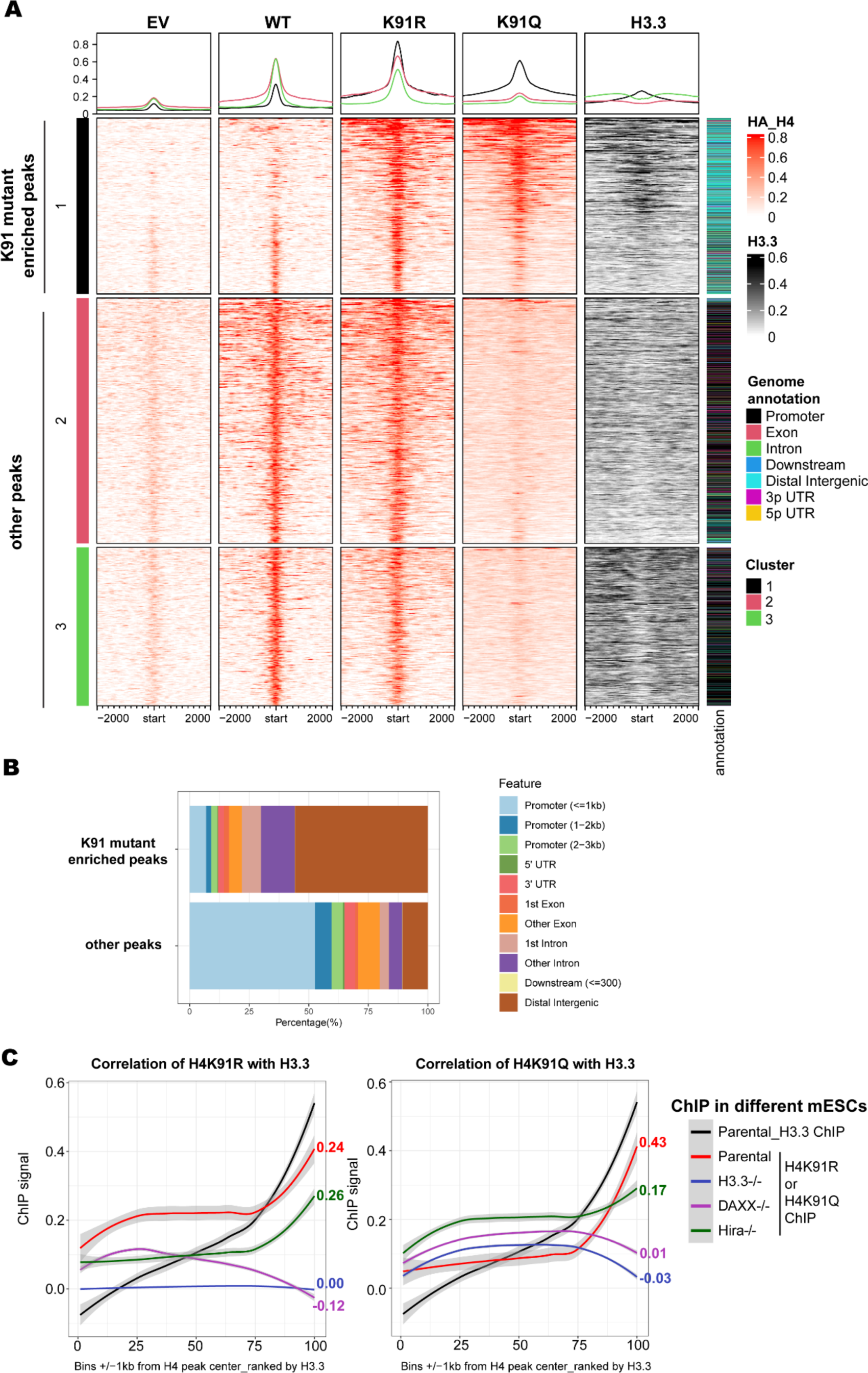
Genome-wide localization of H4K91 mutants in mESCs. Related to Figure 2. (**A**) Heatmap depicting all H4WT, H4K91R, and H4K91Q peaks in mESCs clustered based on HA signal. This signal is quantified in regions centered at H4 peaks and extending ± 2kb. Rows are ranked by mean signal per region in the H4K91R and H4K91Q samples. Cluster 1 is defined as K91 mutant-enriched peaks as shown in Figure 2A, while clusters 2 and 3 are defined as other peaks. The annotation of these peaks with genomic features is shown as a column to the right of the heatmaps. (**B**) Genomic annotation of K91 mutant-enriched peaks (cluster 1) and all other peaks (clusters 2 and 3). (**C**) Correlation of H4K91 mutant ChIP-seq signal (H4K91R: left, and H4K91Q: right) with H3.3 ChIP-seq signal in parental or KO mESCs. Input normalized ChIP-seq signal within bins that comprise regions ±1kb around the center of H4 peaks (union of H4WT, H4K91R, and H4K91Q) are shown. The Pearson’s correlation coefficient (PCC) values was used for this Pearson’s correlation coefficient (PCC) analysis, and PCC values are shown to the right of each figure.

**Figure S3:**
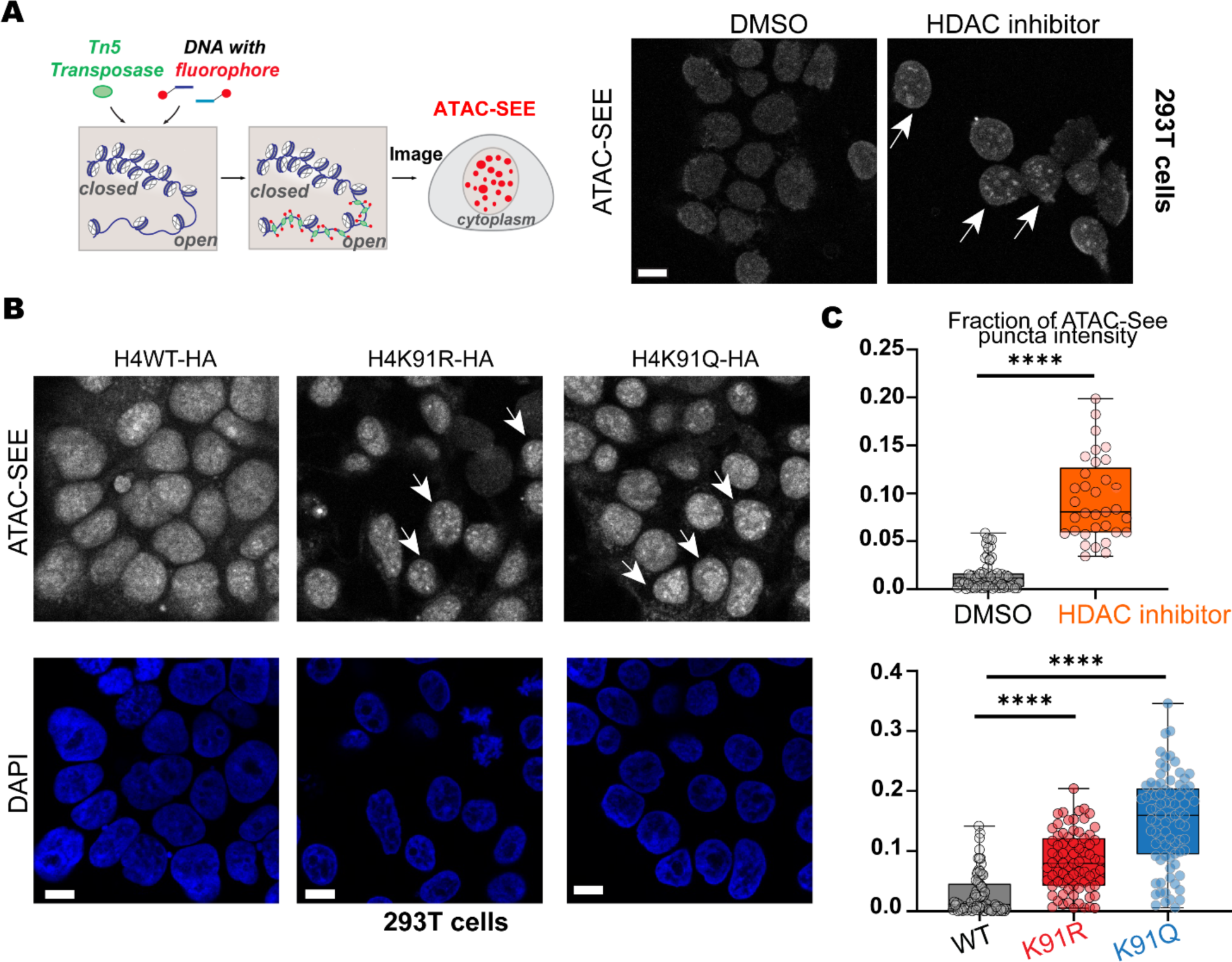
H4K91 mutant expression increases chromatin accessibility. (**A**) Left: Schematic of the assay of transposase-accessible chromatin with visualization (ATAC-see). Right: Representative images of ATAC-see signal in 293T cells treated with DMSO or an HDAC inhibitor (Panobinostat). Arrows indicate cells with distinct ATAC-see signals, which indicate a relatively more open chromatin region. Scale bar: 10μm. (**B**) Representative images of 293T cells expressing WT or mutant H4. DAPI stains the nuclei. Arrows depict cells with distinct ATAC-see signals. Scale bar: 10μm. (**C**) Quantification of in-puncta ATAC-see fluorescence intensity. Fluorescence intensity within the puncta is divided by the whole nucleus intensity. Each dot represents one cell. The box plot shows all data points with the median, and the boundaries indicate the 25^th^ and the 75^th^ percentiles. Statistical significance was calculated using an unpaired t-test. **** *p*<0.0001.

**Figure S4:**
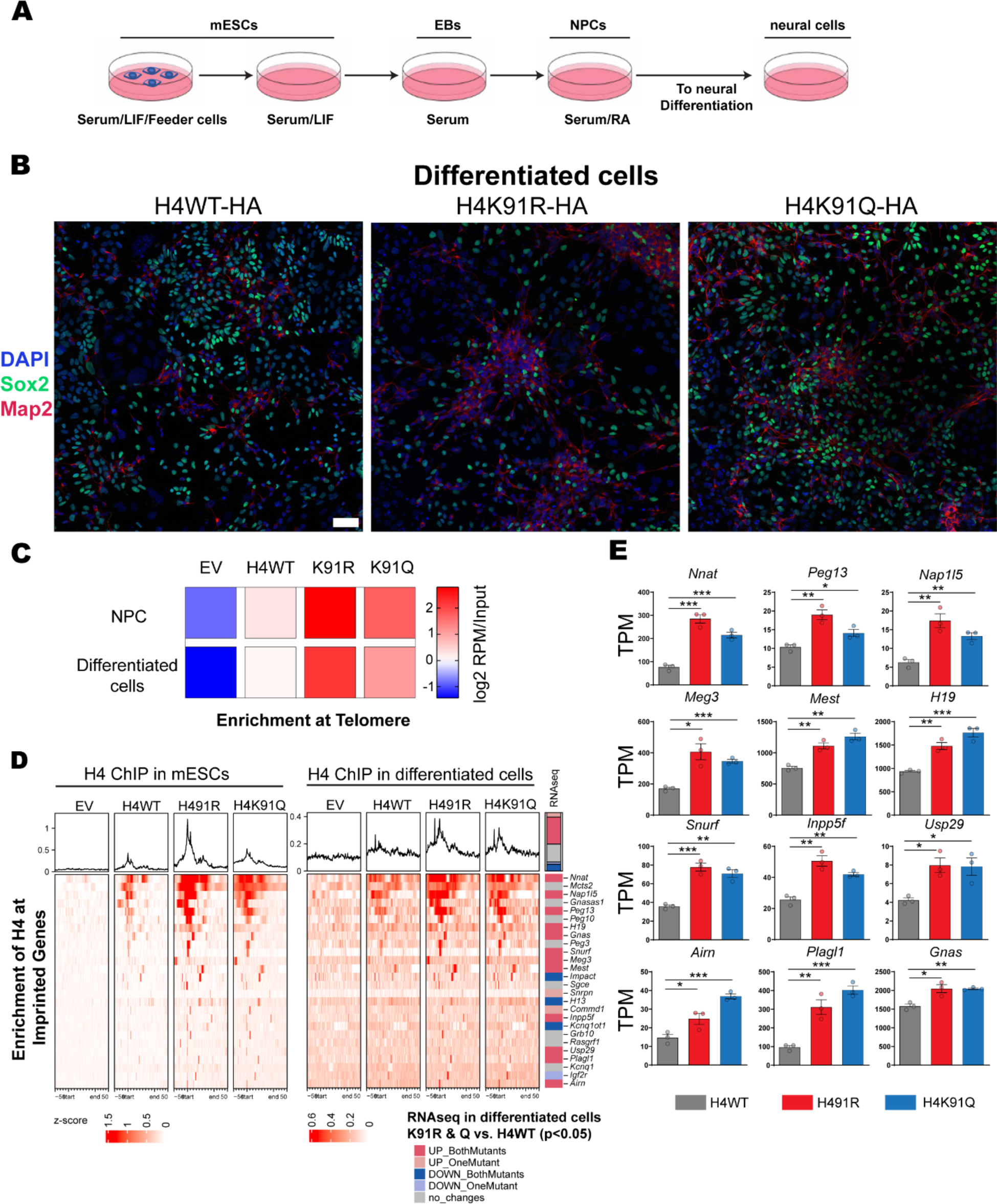
H4 mutants are enriched at telomere and imprinted genes in differentiated neural cells. Related to Figure 4. (**A**) Schematic of mESCs differentiation to embryonic bodies (EBs) and then retinoic acid (RA) mediated differentiation to NPCs, which are further differentiated to neural cells. (**B**) Representative immunofluorescence images of differentiated neural cells 4 days after NPCs were shown. Sox2 stains NPCs, and Map2 labels neurons. Scale bar: 50μm. (**C**) Enrichment of H4K91 mutants at Telomeres. Data are represented in a heatmap of log2 RPM over a matched input. (**D**) Heatmap representation of histone H4 ChIP-seq signal over imprinted genes. The region shown in the heatmap includes the gene body scaled to the same length for each gene, plus 50% of the gene length upstream and downstream. The right column indicates if a gene is significantly changed (*p*<0.05) in differentiated cells expressing H4K91R or H4K91Q compared to H4WT. Rows in the heatmap are sorted by the difference in ChIP signal between the mean of both mutants and WT. (**E**) Visualization of mRNA expression of upregulated imprinted genes. The data represents mean ± s.e.m of TPM values from three independent replicates. Statistical significance was calculated using an unpaired t-test. \**p*<0.05. ** *p*<0.01. *** *p*<0.001.

**Figure S5:**
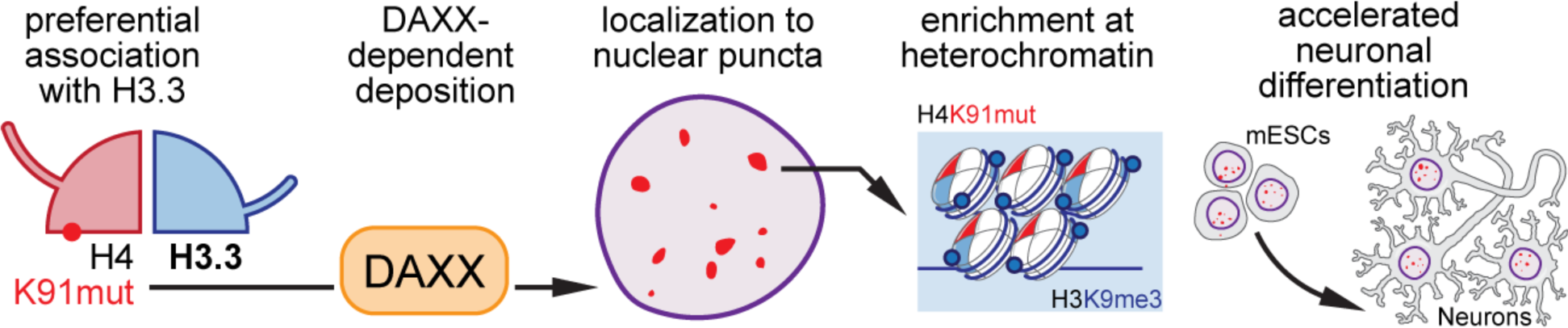
Schematic of the functional roles of histone H4K91 mutants in development. During normal development, wild-type (WT) histone H4 binds to both canonical H3 and histone variant H3.3, therefore having genome-wide incorporation. By contrast, H4K91 mutants prefer binding to histone variant H3.3 and are incorporated into H3.3 and H3K9me3 enriched heterochromatin by a DAXX dependent pathway, which leads to puncta patterns in mESCs as well as differentiated neural cells *in vitro* and *in vivo*. This specific recruitment, combined with the fact that H4K91 mutants destabilize nucleosome stability, leads to more open chromatin, causes aberrant gene expression, and promotes neural differentiation. This derailed cell fate contributes to altered brain development *in vivo.* Future studies will characterize how mutated H4 alters chromatin landscape, gene regulation, and brain development *in vivo* at distinct development times.

## Methods

### ESC culture and differentiation

ESCs are cultured in gelatin-coated plates and serum/LIF medium, which is composed of KO DMEM, 15% ES FBS, β-mercaptoethanol, Glutamax, LIF, Pen/Strep, and non-essential amino acid. To generate ESCs expressing HA-tagged H4 transgenes, H4WT or mutated H4 (H4K91E/R/Q) with 3-HA tagged at the c-terminal were cloned to PiggyBAC transposon plasmid. With the mouse ES Cell Nucleofector Kit (Lonza VPH-1001), PiggyBAC and pBASE plasmid are co-transfected into mESCs, which went through 2 weeks of G418 selection to isolate cells stably expressing H4 transgenes. H3.3 DKO, DAXX KO, and Hira KO mESCs were established previously^31,45,46^. To generate mESCs that transiently express GFP tagged H4 for live cell imaging or staining, H4WT or mutated H4 (H4K91E/R/Q) were cloned to the Xlone-GFP plasmid (addgene 96930), followed by the transfection with Xfect mESC transfection reagent (Takara 631317).

Neural differentiation protocol was modified from the previous method^47^. Briefly, mESCs were cultured with mESC medium in a gelatin-coated dish with MEF feeder cells for two passages, followed by culture without feeder cells for two or three passages. Then 2 million mESCs were plated to LIF-deprived medium and low-attached dishes (Greiner Bio One 633102) for 8 days to form embryonic body (EB) aggregates and 5uM retinoic acid (RA, Millipore R2625) were added to the medium from day 4. The medium was changed every two days. At day 8, EBs were washed and dissociated with fresh Trypsin_EDTA 0.05% (Gibco 25300054) to neuronal progenitor cells (NPCs). NPCs were resuspended in neurobasal medium, which includes neurocult basal medium plus proliferation supplement (stem cell Technologies, 05702), pen/strep, glutamax, EGF (Shenandoah Biotech #100-26) + FGF (Shenandoah Biotech #100-146). Then NPCs were plated on Poly_D_lysine (PDL, Millipore A-003-E) and Mouse laminin (sigma L2020-1MG) coated dishes. The neurobasal medium was changed 2h and one day after plating. Two days after plating, neural differentiation was initiated by switching to the neural differentiation medium (stem cell Technologies, 05704). After partially removing the old medium, a new differentiation medium was added every other day. Cell pellets were collected four days after differentiation for RNA isolation.

### Immunostaining, in situ hybridization, ATAC-see, and Histology

mESCs, NPCs, or differentiated cells were plated on coated glass coverslips (Neuvitro CG-18-PDL). Cells were fixed with 1% fresh paraformaldehyde in PBS for 15-20 minutes and then washed with PBST (0.1% Triton). Then cells were blocked in PBST with 1% BSA (w/v) for 30 minutes before incubating with primary antibodies overnight at 4 ℃, followed by secondary antibody staining at room temperature (RT) for 1h.

mESCs-derived mice at p0 were fixed with 2% fresh paraformaldehyde. After dissection, brain tissues were embedded in parafilm and prepared as 5μm sections by HistoWiz, Inc. To obtain similar cerebral cortex regions among different brain samples, sections were monitored by H&E staining. After rehydrating, brain sections were boiled in sodium citrate buffer (10mM sodium citrate, 0.05% Tween 20, pH 6.0) for 18 min, and then stained with diluted primary antibodies overnight at 4 ℃and then secondary antibody at RT for 1h.

In situ hybridization (ISH) with the telomere probe Tamra-TelG (Tam-OO-TTAGGGTTAGGGTTAGGG 3’, synthesized from BioSynthesis, a gift from Dr. de Lange) was done following de Lange lab published protocols (https://delangelab.org/protocols). Briefly, cells on coverslips were fixed for 5 minutes at RT in 2% paraformaldehyde again. After a wash in PBS, cells were dehydrated in ethanol, consecutively 70%, 95%, and 100% EtOH for 5 minutes each. Next, dry the coverslip and apply a drop of the hybridizing solution including a diluted telomeric probe (1:3000 dilution). Then coverslips with the hybridizing solution were incubated for 3-10 minutes at 70-80 ℃on heatblock, followed by incubation at RT in a humidity-controlled environment.

ISH with brain sections were done using commercially available RNA probes: Tbr2 (ACD, #429641-C2) and Satb2 (ACD, 413261-C3), following their user manual of RNAscope Multiplex Fluorescent Reagent Kit v2 assay.

ATAC-see in 293T cells was done with the EZ-Tn5™ Transposase (Lucigen TNP9211), following the published protocol^36^. Panobinostat (Cayman Chemicals 13280) was used to treat the 293T cells at the concentration of 200nM for 17h.

Imaging of all cells or tissues was done with a 40X or 63X objective and tile scan of Zeiss LSM 780 confocal microscope. Imaging analysis was performed using Zeiss Zen or image J software. Quantification of mutant puncta and ATAC-see puncta were done following the previous published protocol^48^. Following antibodies and dilutions were used: HA (Biolegend 901501, 1:400), ATRX (Santa Cruz, sc-15408, 1:200), DAXX (Santa Cruz, sc-8043, 1:200), Goat anti-Rabbit Alexa Fluor 488/568/647 (Invitrogen, 1:1000), DAPI (1:1000).

### Immunoblot analysis

Cell pellets were resuspended in 1X Laemmli Sample Buffer and boiled for 10-15 minutes at 95℃. Samples were loaded to Tris-Glycine SDS-PAGE gels (16% or 4-20% gels, Invitrogen) and transferred to a nitrocellulose membrane. The blots were first stained with direct blue to detect all proteins, blocked with 1% milk in 1X TBST, and then incubated with diluted primary antibodies overnight at 4℃. After washing with TBST, secondary antibodies were then applied to blots for 1h at RT. Stained immunoblots were incubated in immobilon ECL solution (Millipore) and imaged using an Amersham Imager 600 (GE). Following antibodies were used: HA (Biolegend 901501, 1:1000), H3 (Abcam 1791, 1:10,000), H3.3 (Millipore 09-838, 1:1000), H4 (Abcam ab10158, 1:5000).

### Mononucleosome immunoprecipitation (Mono-IP)

3HA-tagged H4WT, H4K91R, or H4K91Q were cloned into pCDH-EF1-MCS-Puro lentiviral vectors. HEK293T cells that stably express HA tagged H4WT or H4K91 mutants were generated as previously described^10^. HEK293T cells were collected from two 100% confluent 15cm dishes for each immunoprecipitation. Cells were washed with PBS and lysed on ice for 5 minutes in the following buffer: 10 mM HEPES pH 7.9, 10 mM KCl, 1.5 mM MgCl2, 0.34 M sucrose, 10% glycerol, 1mM DTT, protease inhibitor cocktail and Triton X-100 to 0.1%. Then cells were centrifuged at 3.5 K and 4℃for 5 minutes to collect nuclei. Nuclei pellets were washed in lysis buffer without Triton and then resuspended in no-salt solution (3 mM EDTA, 0.2 mM EGTA, 1 mM DTT, and 1× complete protease inhibitor cocktail) on ice with occasional vertexing to break the nuclei. Then microcentrifuge at 4K and 4℃for 5 minutes was done to collect chromatin fraction. Next chromatin pellet was resuspended in MNase digestion buffer (50mM HEPES, 2mM CaCl2, 0.2% NP-40, and 1× complete protease inhibitor cocktail). 18ul MNase (Worthington) was added to digest chromatin at 37℃for 10 minutes and the reaction was stopped by adding EDTA with the final concentration at 5mM. The digested chromatin was centrifuged at max speed for 10 minutes and the supernatant was collected. Take 5% of the supernatant and DNA was extracted by phenol-chloroform after RNase and Proteinase K treatment at 65℃for 1h. The fragment size of DNA was analyzed by 2% agarose gel electrophoresis to see if most nucleosomes were digested into mononucleosomes.

For the next step, digested mononucleosome was dialyzed with M.W. 3500 membrane (Thermo fisher 66330) against the dialysis buffer (20 mM HEPES pH 7.9, 20% glycerol, 0.2 mM EDTA, 0.2% TritonX-100, freshly added protease inhibitors) with 150mM KCl for 2h at 4℃. Then dialyzed mononucleosome was incubated with 100 ul anti-HA magnetic beads (Thermo 88837) overnight at 4℃while rotating. Then beads were washed three times using dialysis buffer with 150mM KCl and then three times through dialysis buffer with 100mM NaCl and 10% glycerol. Elution was done using a 5% acetic acid solution. Three replicates of monoIP pulldown were collected and analyzed by mass spectrometry analysis and immunoblot analysis.

### Mass spectrometry

MonoIPs were dried in a centrifuge vacuum concentrator and resuspended in 20 ul of 50mM Ammonium Bicarbonate. Samples were propionylation as previously described^49^. In short, 10ul of propionic anhydride was added to 30ul of Acetonitrile and 10ul of the mixture was added to each sample and well mixed. Immediately, the pH of each sample was adjusted by addition of ammonium hydroxide to pH 8.0. Samples were incubated for 15 min at room temperature, dried down, and the propionylation was repeated. Samples were resuspended in 50mM Ammonium Bicarbonate and digested overnight with 2 ul of trypsin at room temperature. Digested peptides were dried down and the propionylation steps were repeated. Samples were desalted with self-packed C18 stage-tips and resuspended in 20ul 0.1% formic acid. The samples (2ul) were injected onto the Acclaim PepMap 100 C18 column (3 µm x 0.075 mm x 150 mm) for the analysis on a Q-Exactive Plus instrument (Thermo Fisher Scientific) attached to an UltiMate 3000 UHPLC (Thermo Fisher Scientific) and Nanospray Flex ion source (Thermo Fisher Scientific). The peptides were separated using buffer A (0.1% formic acid) and buffer B (80% acetonitrile and 0.1% formic acid) with a gradient of 2-35% over 50 min. The column was then washed at 98% buffer B over 5 min and equilibrated to 2% buffer B. Data-independent acquisition (DIA) was performed with the following settings: A full MS1 scan from 300 to 1100 m/z with a resolution of 70,000, an automatic gain control (AGC) target of 1 × 106, and a maximum injection time of 50 ms. DIA scans were attained across the same mass range with sequential isolation windows of 24 m/z with a normalized collision energy of 30, a resolution of 17,500, an AGC target of 2 × 105, and a maximum injection time of 60 ms. Data was analyzed with in-house program, EpiProfile.

### Chromatin immunoprecipitation (ChIP)

For each immunoprecipitation, mESCs from two confluent T75 flasks were fixed with 1% paraformaldehyde for 5 minutes with gentle rotation and then quenched with 125 mM glycine for 5 minutes. To get chromatin, fixed cells were resuspended in 1ml of buffer 1 (50 mmol/L HEPES pH 7.5, 140 mmol/L NaCl, 1 mmol/L EDTA, 10% glycerol, 0.5% NP-40, 0.25% Triton X-100, and 1x complete protease inhibitor cocktail) and incubated for 10 minutes at 4℃with rotation. Samples were centrifuged at 1400x g for 5 minutes and pellets were resuspended in 1ml of buffer 2 (10 mmol/L Tris-HCl pH 8.0, 200 mmol/L NaCl, 1 mmol/L EDTA, 0.5 mmol/L EGTA, and 1× complete protease inhibitor cocktail) for incubation at 4℃for 10 minutes with rotation. Next samples were centrifuged at 1400xg for 5 minutes and resuspended in 900ul of buffer 3 (10 mmol/L Tris-HCl pH 8.0, 100mmol/L NaCl, 1 mmol/L EDTA, 0.5 mmol/L EGTA, 0.1% sodium deoxycholate, 0.5% N-Lauroylsarcosine, and 1× complete protease inhibitor cocktail). Samples were homogenized by passing through a syringe (28G1/2) eight times. Finally, 100ul of 10% Triton X-100 was added to homogenized chromatin for 16 minutes of sonication using a Covaris E220.

Sonicated samples were centrifuged for 10 minutes at 4℃. 5% of the supernatant was saved to extract DNA as input, while the rest was incubated with 75ul anti-HA magnetic beads (Thermo 88837) overnight at 4℃while rotating. Beads were washed six times with 1ml of cold RIPA buffer (50 mmol/L HEPES pH 7.5, 500 mmol/L LiCl, 1 mmol/L EDTA, 0.7% sodium deoxycholate, and 1% NP-40) and once with 1ml of wash buffer (10 mmol/L Tris-HCl pH 8.0, 50 mmol/L NaCl, and 1 mmol/L EDTA). Then beads were incubated with 210 ul of elution buffer (50 mmol/L Tris-HCl pH 8.0, 10 mmol/L EDTA, and 1% SDS) at 65℃for 3 minutes while shaking. The eluted chromatin was centrifuged at maximum speed for 1 minute at RT. Cross-linking was reversed by incubating the supernatant overnight at 65℃, followed by 1h RNaseA treatment at 37℃and 2h Proteinase K treatment at 55℃. DNA was extracted after mixing with phenol-chloroform and then ethanol precipitation.

For anti-HA ChIP with NPCs or differentiated neural cells, ten million cells were collected and fixed with 1% paraformaldehyde for 5 minutes with gentle rotation and then quenched with 125 mM glycine for 5 minutes. To get chromatin, fixed cells were washed with cold 1X PBS and lysed using previously published NEXSON protocol^50^. Briefly, cells were resuspended in 500ul of lysis buffer (5 mM PIPES, pH 8, 85 mM KCl, 0.5% NP-40, 1X complete Protease inhibitor cocktail) and sonicated in Bioruptor Plus (low power, 9 cycles of 15 seconds on and 30 seconds off). A small fraction of sonicated samples were checked under the microscope in between to monitor the lysis efficiency until more than 70% of cells were lysed. Then lysed samples were centrifuged for 5 minutes at 1000xg at 4 ℃. Nuclei pellets were washed in lysis buffer and resuspended in 130ul sonication buffer (10 mM Tris-HCl, pH 8, 0.1% SDS, 1 mM EDTA, 1X Complete Protease inhibitor cocktail) for 6min sonication using a Covaris E220. The sheared chromatin was incubated with 4ul HA antibody (Biolegend 901501, 1:1000) and immunoprecipitation was done using ChIP-IT high-sensitive kit (active motif 53040).

ChIP libraries were generated with input and pulldown DNA using NEBNext Ultra II DNA Library Prep Kit for Illumina (NEB E7645L) for sequencing on an Illumina Nextseq with 75-bp read length and single-end.

### RNA isolation and sequencing

Total RNA was isolated from cells using the RNeasy kit (Qiagen) and analyzed using the Bioanalyzer RNA Pico (Agilent) before library preparation. PolyA mRNA enrichment was performed using the NEB Next Poly(A) mRNA magnetic isolation module (NEB E7490L), followed by library preparation using the NEB Next Ultra II RNA Library Prep Kit (NEB E7770L). Sequencing was done using the Illumina Nextseq with 75-bp read length and single-end.

### Bioinformatic analysis

#### RNA-seq analysis

Transcript abundance was determined from FASTQ files using Salmon (v0.8.1) and the GENCODE reference transcript sequences^51^. Transcript counts were imported into R with the tximport R Bioconductor package (v1.8.0), and differential gene expression was performed with the DESeq2 R Bioconductor package (v1.20.0)^52,53^. Normalized counts were retrieved from the DESeq2 results and z-scores for the indicated gene sets were visualized with heatmaps generated using the pheatmap R package (v1.0.12)^54^. Gene ontology analysis was performed with the selected gene lists using the clusterProfiler R Bioconductor package (v4.0.5)^55^.

#### ChIP-seq analysis

ChIP-seq reads were aligned using the Rsubread R Bioconductor package (v1.30.6) and predicted fragment lengths were calculated by the ChIPQC R Bioconductor package (v1.16.2)^56,57^. Normalized, fragment-extended signal bigWigs were created using the rtracklayer R Bioconductor package (v1.40.6), and peaks were called using MACS2 (v2.1.1)^58,59^. Range-based heatmaps showing signal over genomic regions were generated using the profileplyr R Bioconductor package (v1.8.1)^60^. To identify K91mut enriched peaks, the union of HA peaks from H4WT, H4K91R, and H4K91Q cells was filtered to those with a p value less than 10^-10 (as reported from MACS2), and then peaks from empty vector cells were removed. K-means clustering was used on the regions around these peaks to cluster the signal heatmaps and identify a cluster with high signal in both H4 mutant samples and low signal in H4WT expressing cells. Peaks were annotated with the various types of genomic regions using the ChIPseeker R Bioconductor package (v1.28.3)^61^. Any regions included in the ENCODE blacklisted regions of the genome were excluded from all region-specific analyses^62^. To assess ChIP-seq signal in telomeres, fastq files were aligned to a DNA sequence of 150 conserved telomere repeats (TTAGGG) using the R Bioconductor package Rbowtie2 (v1.12.0)^63,64^.

#### Genome-wide correlation analysis

The signal from ChIP samples was quantified in 20bp bins +/− 1kb from the center of the HA union peak set described above. These bins were then ranked by the indicated ChIP sample, and the bins were then divided into 100 quantiles for the x-axis of the correlation plot. The signal of all ChIPs in these ranked quantiles was then plotted using the geom_smooth() function from the ggplot2 R package (v3.3.6)^65^.

### Tetraploid complementation and all-ESC mutant mice

Animals were housed and cared for according to a protocol approved by the IACUC of Weill Cornell Medical College (Protocol number: <u>2014-0061</u>). Wild-type ICR mice were purchased from Taconic Farms (Germantown, NY). Females were super-ovulated at 6–8 weeks with 0.1 ml CARD HyperOva (Cosmo Bio Co., Cat. No. KYD-010-EX) and 5 IU hCG (Human chorionic gonadotrophin, Sigma-Aldrich) at intervals of 48 hours. The females were mated individually to males and checked for the presence of a vaginal plug the following morning. Plugged females were sacrificed at 1.5 days post-hCG injection to collect 2-cell embryos. Embryos were flushed from the oviducts with advanced KSOM (Cat # MR-101-D, Millipore). The 2-cell embryos were subjected to electrofusion to induce tetraploidy. Fused embryos were moved to new KSOM micro drops covered with mineral oil and cultured to the blastocyst stage until ESC injection.

ESCs expressing HA-tagged H4WT, H4K91R, and H4K91Q were trypsinized, resuspended in ESC medium, and kept on ice. A flat tip microinjection pipette was used for ESC injections. ESCs were collected at the end of the injection pipette and 10–15 cells were injected into each 4n blastocyst. The injected 4n blastocysts were kept in KSOM until embryo transfer. Typically, ten injected 4n blastocysts were transferred into each uterine horn of 2.5 dpc pseudo-pregnant ICR females. After injecting ESCs into tetraploid blastocysts, the 4n cells of the host embryo contribute solely to the placenta, while the injected ESCs form the embryo properly. All-ESC pups were recovered via cesarean section at embryonic day 19.5 (E19.5), which is equivalent to postnatal day 0 (P0) in normal fertilized embryos Mice genotyping was done using PCR with the following two primers:

Forward: 5’ CGGACTAGTGCCACCATGTCTGG 3’

Reverse: 5’ AAATATGCGGCCGCTTAAGCATAATCAGGCACAT 3’

